# Biocosm: A Simulation Environment for Designing Energy-Constrained Ethological Sensors

**DOI:** 10.64898/2026.05.18.725991

**Authors:** Matt Gaidica

**Affiliations:** Department of Neuroscience, Washington University in St. Louis

**Keywords:** measurement-process simulation, ethological sensing, proximity logging, energy-constrained instrumentation, agent-based simulation

## Abstract

Long-duration ethological sensing increasingly depends on miniaturized, battery-constrained devices that must decide when to observe, transmit, store, compute, or sleep. These policy decisions shape not only device lifetime, but also which biological events are ultimately available for inference. This manuscript presents Biocosm, a measurement-process simulation framework for energy-constrained ethological instrumentation. Biocosm separates the latent biological world from the observation model, sensing policy, energy model, and analysis layer, allowing researchers to evaluate how alternative sensing strategies transform true behavioral interactions into observed data. Although motivated by wireless proximity logging, the framework is hardware-agnostic: beaconing, listening, sensing, storage, and computation are represented as parameterized state/action costs rather than as fixed properties of one device architecture. This structure supports reproducible design-space exploration, policy comparison, and sensitivity analysis before field deployment. Biocosm is grounded in prior work on animal proximity loggers, low-power wireless discovery and energy modeling, agent-based ethological simulation, and digital twin-inspired methods in medicine and neurotechnology. Framing measurement as a coupled world–observation–policy system makes instrumentation choices explicit and quantifiable.

## 1 Introduction

Automated ethological sensing has expanded the temporal and spatial scale at which animal behavior can be measured. Animal-borne proximity loggers, biologgers, radio-frequency identification systems, wireless sensor networks, and machine-vision pipelines now allow researchers to study social contact, movement, activity, and physiology across conditions that would be difficult or impossible to observe manually (Drewe JA et al. 2012; Kirkpatrick et al. 2021; Levin II et al. 2015; Prange S et al. 2006; Ripperger SP, Carter GG, et al. 2020; Ripperger SP, Josic D, et al. 2016; Rutz C et al. 2015; Ryder TB et al. 2012; Smith JE and Pinter-Wollman N 2021). These tools are especially valuable for studying nocturnal, cryptic, socially complex, or long-duration behaviors, where direct observation is sparse or disruptive.

However, ethological sensors are not passive windows into biological reality. Recent animal social-sensing work shows that sampling frequency, sampling duration, sampling coverage, and measurement method can alter inferred interactions and network structure (Gelardi V et al. 2020; He P et al. 2023). In practice, every deployment encodes an observation policy. A device may sample continuously, sleep between bursts, beacon periodically, listen for nearby nodes, buffer events locally, transmit opportunistically, or adapt its sampling schedule based on recent observations. Each of these choices affects energy consumption and the probability that a true behavioral event is detected. In small-animal systems, the trade space is especially constrained: battery capacity, device mass, memory, antenna geometry, and animal welfare all limit how aggressively a device can observe the world (Huels F 2025; Kirkpatrick et al. 2021; Levin II et al. 2015; Walker AM 2024).

This creates a methodological problem. The scientific data produced by an ethological sensing system are jointly determined by the animal, the environment, the sensing modality, the embedded firmware policy, and the energy budget. A missed contact may reflect biological absence, poor radio propagation, insufficient sampling duty cycle, body shielding, depleted battery, device-specific variation, stochastic false negatives, or post-processing choices (Boyland NK et al. 2013; Ossi F 2016). Prior validation studies have repeatedly shown that proximity-logger outputs depend on distance, orientation, habitat, attachment geometry, inter-device variation, signal thresholds, and false-negative detection processes (Boyland NK et al. 2013; Camargo VA 2025; Drewe JA et al. 2012; Kirkpatrick et al. 2021; Ossi F 2016; Prange S et al. 2006; Rutz C et al. 2015; Ryder TB et al. 2012). Wireless systems literature similarly shows that discovery probability, latency, and energy use are coupled through advertising, scanning, transmission, and sleep schedules (Cho K et al. 2015; Dutta P and Culler D 2008; Kindt PH et al. 2020; Liu J, Chen C, and Ma Y 2012; Luo B and Liu J 2014; Siekkinen M et al. 2012; Sun W et al. 2014).

Biocosm addresses this problem by treating the sensing system as part of the scientific method. Rather than asking only whether a given device works, Biocosm asks a broader question: given a plausible biological world and a candidate sensing policy, what data would this instrument produce, at what energy cost, and with what measurement bias? This reframes ethological instrumentation as a coupled world–observation–policy–energy–analysis pipeline.

## 2 Conceptual Framework: A Digital Twin-Inspired Measurement Model

This manuscript uses **digital twin-inspired** language in a focused methodological sense. Biocosm is not a full biophysical animal model, a full electromagnetic channel simulation, or a device-specific firmware emulator. Instead, it is an executable representation of the coupled measurement process formed by animals, sensors, observation rules, communication constraints, and energy budgets.

This framing is inspired by broader digital twin work in medicine and neurotechnology, where computational models are increasingly used to link latent biological state, observation models, and intervention planning (Corral-Acero J 2020; Popescu D 2024; Wang HE 2024). It also follows general digital-twin formulations that define a twin as a virtual representation of a physical object or process updated through data flows from the physical system (Al-Ali AR et al. 2020; Khan LU et al. 2022; Minerva R, Lee GM, and Crespi N 2020). In virtual brain twin frameworks, for example, generative neural dynamics are conceptually separated from observation models and intervention models (Wang HE 2024). Biocosm applies a related separation to ethological instrumentation: latent animal behavior is distinct from what sensors can observe, and sensor policy is distinct from the biological process being measured. The current implementation is an offline, scenario-based executable measurement model rather than a continuously synchronized twin of a live deployment.

Biocosm is organized around five conceptual layers:

1. **World layer** — the latent biological world, including animals, movement along paths and nodes as representative routes by which behavior is constrained, social proximity, circadian state, arena geometry, and true interaction events.
2. **Signal or observation layer** — the transformation from latent events into observable evidence, such as proximity signal strength, detection probability, sensing noise, or event fragmentation.
3. **Policy layer** — the embedded rules that schedule sleep, wake, sample, listen, and beacon states (and adaptive transitions among them), including rules that reduce or alter sampling during inactivity or within scheduled rest/sleep windows.
4. **Energy layer** — the cost of each state or action, expressed as a configurable energy model rather than a fixed hardware assumption.
5. **Analysis layer** — the metrics applied to observed logs, including event capture, efficiency, inferred social networks, observation bias, and policy frontiers.

This layered decomposition (Figure 1) is consistent with digital-twin architectures in IoT and wireless systems that separate physical interaction, communication, virtual or twin representation, analytics, and application functions (Al-Ali AR et al. 2020; Khan LU et al. 2022).

**Figure 1:**
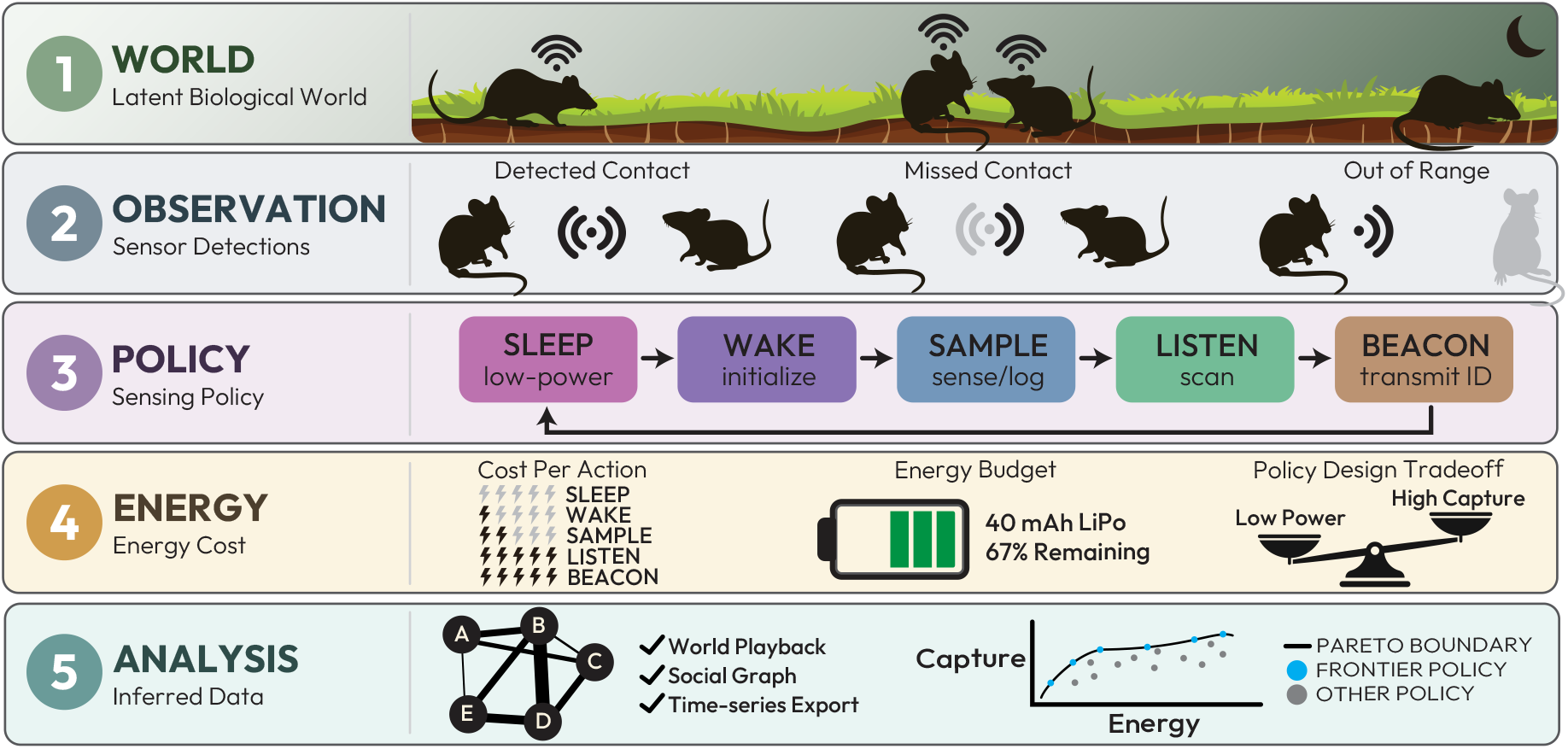
Biocosm conceptual architecture. The framework separates the latent biological world, imperfect observation, sensing policy, energy cost, and downstream analysis. World and observation layers define ground truth and detectable evidence; the policy layer executes duty-cycled sense/listen/beacon actions under an energy budget; the analysis layer evaluates observed logs relative to ground truth, including social-network inference and capture–energy trade-offs.

The key methodological separation is that the policy layer is not allowed to access ground truth. Simulated sensing nodes operate only on observable history, while the simulator retains ground truth for later evaluation. This distinction allows the same biological world to be replayed under multiple candidate policies, making it possible to attribute differences in observed data to measurement strategy rather than uncontrolled biological variation.

## 3 Related Work

### 3.1 Animal proximity logging and validation

Animal proximity loggers have a long validation tradition. Early wildlife radiocollars demonstrated that automated proximity detection could quantify contacts among animals that are difficult to observe directly, while also showing that detection distance, antenna orientation, and field conditions require explicit validation (Prange S et al. 2006). Drewe and colleagues validated proximity loggers for cattle and badgers and found that logger identity could be recorded accurately, but contact duration could be fragmented, motivating post-processing rules such as amalgamation windows (Drewe JA et al. 2012).

Encounternet studies extended proximity logging to smaller animals and more complex social-network questions. Ryder and colleagues showed that proximity loggers can increase both the quantity and quality of social-network data in wild birds (Ryder TB et al. 2012). Rutz and colleagues provided a particularly important methods precedent by combining field experiments, theoretical modeling, statistical modeling, behavioral observations, and computer simulation to calibrate animal-borne proximity loggers (Rutz C et al. 2015). Levin and colleagues further demonstrated field use of miniaturized Encounternet tags in small birds, while cautioning that RSSI-thresholded networks are more defensible than exact distance estimates (Levin II et al. 2015).

Small-animal biologging systems have continued to advance. Ripperger and colleagues developed lightweight sensor nodes for bats and later a wireless biologging network that enabled direct proximity sensing, high-resolution tracking, and remote data download in small vertebrates (Ripperger SP, Carter GG, et al. 2020; Ripperger SP, Josic D, et al. 2016). That work is especially relevant because it treats energy modeling and adaptive operation as central design considerations rather than implementation details (Ripperger SP, Carter GG, et al. 2020).

Recent BLE-oriented systems further motivate a general simulation framework. ProxLogs introduced miniaturized BLE proximity loggers for small animals (Kirkpatrick et al. 2021).

Subsequent field work using miniaturized BLE proximity loggers in agricultural settings showed that scan interval, habitat, logger height, and device variation affected battery life and signal strength, requiring context-specific calibration (Huels F 2025). BLE systems for grazing sheep similarly demonstrated the potential of low-power wireless proximity and localization, while showing that range and localization accuracy depend on device placement and animal behavior (Walker AM 2024).

Across these studies, a recurring lesson is that proximity data are not ground truth. They are conditional observations shaped by hardware, firmware, environment, inter-logger variation, false-negative detections, and post-processing (Boyland NK et al. 2013; Drewe JA et al. 2012; Huels F 2025; Kirkpatrick et al. 2021; Levin II et al. 2015; Ossi F 2016; Rutz C et al. 2015). Biocosm formalizes this lesson by making the observation process executable and inspectable.

### 3.2 Radio discovery, sensing schedules, and energy trade-offs

Low-power wireless systems are governed by coupled trade-offs among discovery probability, latency, bandwidth, and energy. BLE is a clear example because neighbor discovery depends on temporal overlap between advertising events and scanning windows (Cho K et al. 2015; Liu J, Chen C, and Ma Y 2012). More generally, asynchronous neighbor-discovery theory shows that low duty cycle, bounded latency, and energy efficiency are coupled even outside BLE-specific implementations (Dutta P and Culler D 2008; Sun W et al. 2014). Analytical models of BLE discovery describe how advertising interval, scan interval, and scan window affect discovery probability and expected latency (Cho K et al. 2015; Liu J, Chen C, and Ma Y 2012). Energy models extend this trade space by accounting for the current consumed in advertising, scanning, receiving, transmitting, sleeping, and state transitions (Kindt PH et al. 2020; Luo B and Liu J 2014; Siekkinen M et al. 2012).

This literature supports Biocosm ’s treatment of policy as a first-class experimental variable. A sensing node’s behavior is not defined only by its sensor or radio; it is defined by when that sensor or radio is active. Sparse listening may extend lifetime but miss brief events. Frequent beaconing may increase observability but consume energy or create channel contention. Adaptive policies may improve efficiency by sampling more aggressively after recent detections, but they may also create behavior-dependent sampling bias.

Animal studies have reached similar conclusions empirically. BLE proximity studies report that scan interval affects battery life and detection performance, and that habitat, logger height, device placement, and animal behavior shape the relationship between signal strength and inferred proximity (Huels F 2025; Walker AM 2024). Livestock proximity-logger validation studies further show that high specificity and precision can coexist with low sensitivity, making false negatives a central design consideration (Camargo VA 2025). GPS-based studies at wildlife– livestock interfaces illustrate the same measurement-design issue at the analysis level: inferred interaction frequencies depend on explicit spatial and temporal encounter definitions (Triguero-Ocaña R et al. 2019). These findings motivate a general, parameterized energy and observation model rather than a single fixed hardware profile.

### 3.3 Biological simulation and digital twins

Simulation has become a methodological tool for designing biological measurement systems, not only for explaining completed experiments. In animal social-network analysis, agent-based modeling has been used to evaluate sampling design before empirical deployment (Kaur R 2025). Recent work in animal-society sensing has also used data-informed subsampling and robustness analyses to evaluate how study-design choices affect inferred networks before or alongside deployment (Gelardi V et al. 2020; He P et al. 2023). Movement simulation packages such as `abmAnimalMovement` reflect growing interest in accessible agent-based methods for ecological and ethological research (Fournier AMV 2022).

More broadly, biological simulation platforms such as BioDynaMo and Morpheus show that modular, reproducible simulation environments are themselves publishable scientific infrastructure (Breitwieser L 2021; Starruß J et al. 2014). These platforms are not proximity-logging tools, but they demonstrate how biological simulation can separate mechanism, parameters, execution, and analysis in a reusable framework.

Digital twin approaches in medicine and neuroscience provide an additional conceptual precedent. Precision cardiology digital twin frameworks argue that individualized computational models can support diagnosis, prediction, and intervention planning (Corral-Acero J 2020). Scoping reviews in digital health describe digital twins as an emerging research-and-development landscape spanning organ, system, and whole-body modeling (Popescu D 2024). Virtual brain twin frameworks similarly emphasize personalized, generative, adaptive models linked to observation and intervention (Wang HE 2024). Biocosm adapts this digital twin logic to ethological instrumentation: the goal is not to replace empirical validation, but to provide a reproducible model of how measurement choices transform biological events into analyzable data.

## 4 Biocosm Framework

### 4.1 Overview

Biocosm is a simulation framework for energy-constrained ethological sensing. It is designed to evaluate sensing policies under controlled, reproducible biological scenarios. The framework simulates a latent world, generates true events, applies an observation model, executes sensing policies, accumulates energy cost, and exports observed logs and performance metrics.

The central abstraction, illustrated in Figure 1, is:

latent biological world → imperfect observation → sensing policy → energy cost → observed data → scientific inference

This abstraction intentionally avoids tying the method to one device architecture. A wireless proximity logger, acoustic recorder, camera trap, implantable neural device, accelerometer, RFID reader, or environmental sensor may all be represented if the relevant observation and energy models can be parameterized. In the current software implementation (sections 4.3 and 4.8 and Figures 2 and 3), a reference wireless beacon/listener workflow—informed by prior BLE wearable development (Gaidica and Dantzer 2022)—illustrates one instantiation; duty-cycle timings, detection parameters, and per-state energy costs are user-specified rather than fixed by the framework.

**Figure 2:**
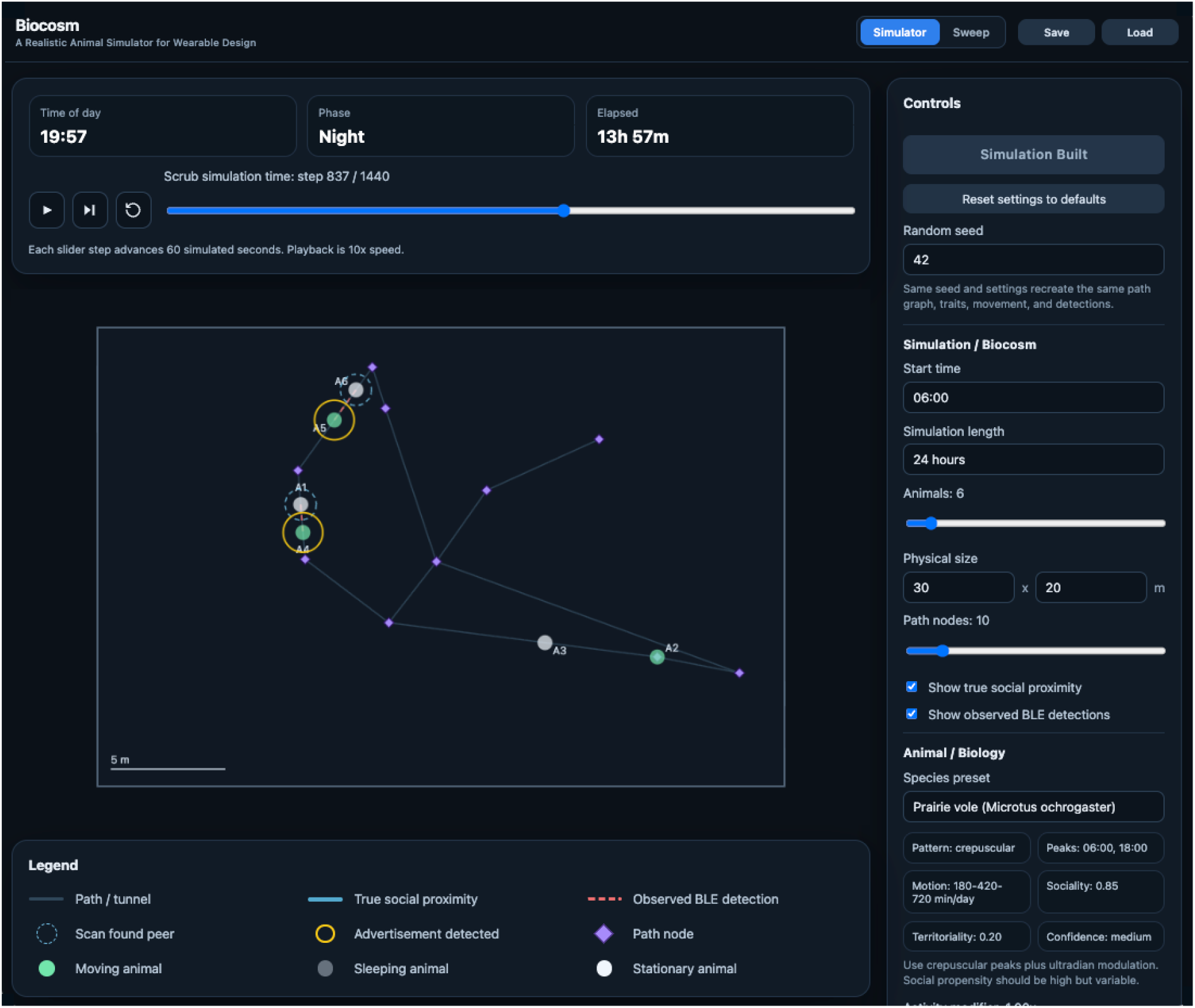
Primary Biocosm user interface (Simulator workspace). Users configure species presets, enclosure geometry, policy parameters, and energy models; build a seed-deterministic simulation; and inspect ground-truth proximity, observed detections, scan/beacon activity, and time-series metrics on a shared timeline. The timeline can replay the full behavioral sequence and capture events so users can validate simulated movement, proximity, and detection against policy-driven observations before committing to sweeps or deployment. Additional workspaces support batch policy sweeps and surrogate-assisted policy optimization.

**Figure 3:**
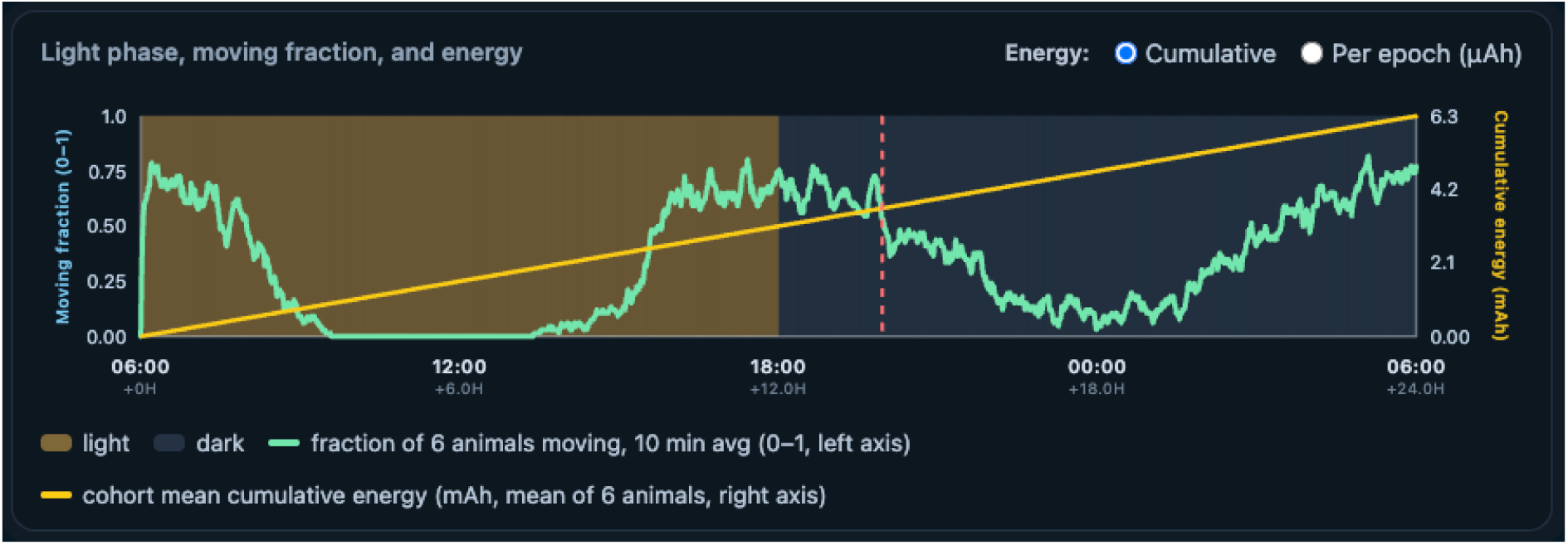
Light phase, moving fraction, and device energy in the Biocosm simulator. Summary panel from a representative run with *n* = 6 prairie voles, showing reproduction of the imposed light cycle, population movement profile, and cumulative energy use over the simulation length. Together with timeline replay (Figure 2), these traces support validation of species presets, circadian structure, and energy accounting before policy sweeps or export.

### 4.2 World layer

The world layer represents the latent biological state. Animals occupy a simulated environment and generate ground-truth proximity or interaction events independent of whether any device observes them. World parameters may include the number of animals, arena dimensions, a path graph of nodes and connecting routes, movement speed, interaction radius, circadian modulation, social preference structure, activity-state transitions, and random seed.

The purpose of the world layer is not to reproduce all biological complexity. Instead, it provides a controlled substrate for testing measurement systems. By holding the world constant across policy comparisons, Biocosm allows researchers to quantify how alternative sensing strategies distort the same underlying biological process.

### 4.3 Species-specific behavior presets

Rather than requiring users to hand-tune circadian windows, bout lengths, and social tendencies, Biocosm provides **species presets**: literature-informed archetypes for laboratory and wild/house mouse (*Mus musculus*), laboratory rat (*Rattus norvegicus*), deer mouse (*Peromyscus maniculatus*), white-footed mouse (*P. leucopus*), prairie vole (*Microtus ochrogaster*), meadow vole (*M. pennsylvanicus*), bank vole (*Myodes glareolus*), fox squirrel (*Sciurus niger*), eastern gray squirrel (*S. carolinensis*), red and ground squirrel archetypes, and a human profile. Each preset specifies an activity pattern class (nocturnal, diurnal, crepuscular, or ultradian), clock-time activity peaks, broad active/rest windows, expected daily motion-positive minutes, movement and rest bout scales, individual phase offsets, and triangular distributions (`min, mode, max`) for social propensity and territoriality. At initialization, the simulator samples one trait vector per animal from these distributions and holds it fixed for the run, preserving individual heterogeneity while keeping population-level priors explicit and exportable.

The presets are simulation priors for *collar-relevant* surface behavior—when animals are likely to move, remain inactive, or co-locate—rather than literal EEG-defined sleep ethograms (Hazlerigg and Tyler 2019). Qualitative structure was compiled from published activity and rest– bout studies. Laboratory rodents are modeled as nocturnal and polyphasic (Pernold, Rullman, and Ulfhake 2023; Simasko and Mukherjee 2009; Stephenson et al. 2013; Wang et al. 2020). Prairie and meadow voles incorporate crepuscular peaks and ultradian activity–rest cycling (Bueno-Junior et al. 2023; Lewis and Curtis 2016; Van Rosmalen and Hut 2021). Tree squirrels use diurnal, often bimodal daylight activity with high territoriality and lower default social attraction (Thompson 1977; Wassmer and Refinetti 2019). Deer mice follow nocturnal, bimodal feeding–activity patterns (Jaeger 1982). Each preset carries a confidence label (`high, medium`, or `low`) to signal how strongly the numerical ranges are supported for simulation purposes; users can apply global modifiers (activity level, sociality, bout length) without overwriting the underlying species profile.

These presets are priors for scenario generation and sensitivity analysis, not validated species models for direct biological inference.

The world engine combines circadian, optional ultradian, social-context, and stochastic components to drive movement along a path graph inside a rectangular enclosure. This design supports a central Biocosm question: under a biologically plausible activity landscape, how do alternative sensing schedules (BLE in the reference implementation) recover proximity structure at acceptable energy cost?

### 4.4 Observation layer

The observation layer converts latent events into detectable evidence. For wireless proximity sensing, this may include distance-dependent signal attenuation, stochastic detection probability, orientation effects, body shielding, habitat attenuation, packet overlap, signal thresholds, and device-to-device variation. For other modalities, the observation layer could represent field of view, acoustic range, accelerometer thresholding, motion-triggered camera probability, or sensor noise.

A generic observation model can be written as:

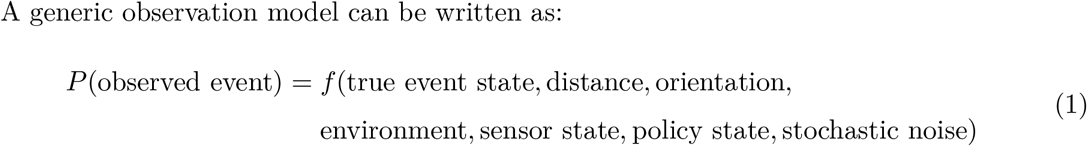

This probabilistic framing is important because many ethological sensing systems are neither perfectly sensitive nor perfectly specific. Observed logs are samples from an imperfect measurement process, not direct copies of ground truth.

### 4.5 Policy layer

The policy layer defines when each sensing node transitions among sleep, wake, sample, listen, and beacon states (Table 1). Policies may be fixed, adaptive, randomized, or state-dependent. Examples include periodic observation, burst sampling, duty-cycled listening, scheduled beaconing, event-triggered high-duty sampling, energy-aware throttling, and adaptive policies based on recent detections. The layer also supports rules that down-regulate or stretch inactive scan and advertise schedules when onboard activity is low or when the collar is within species-informed sleep windows, so duty cycles can track expected rest rather than run at full sampling intensity continuously.

**Table 1:**
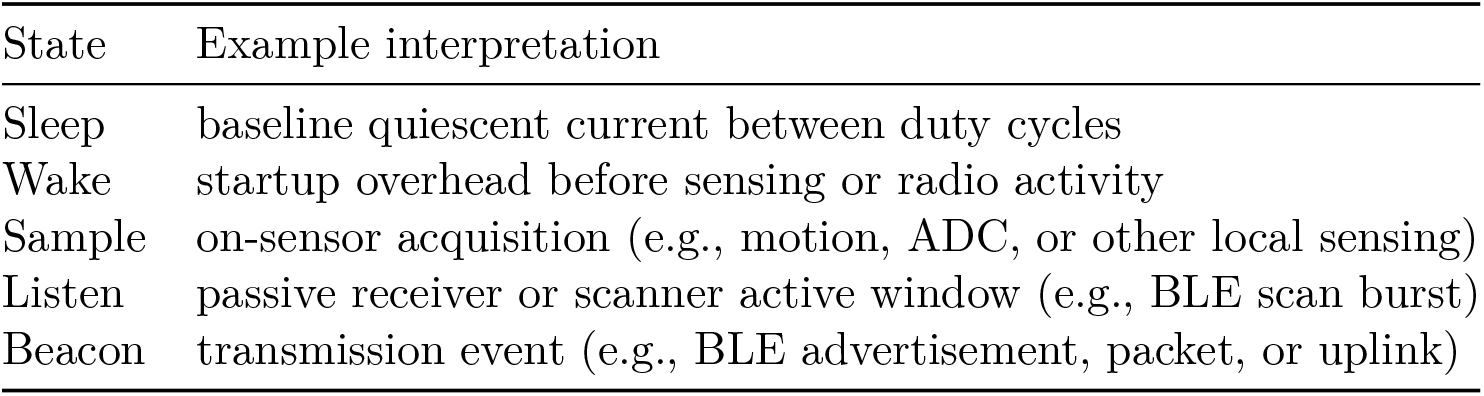
Representative state and action classes in the energy layer.

| State | Example interpretation |
| --- | --- |
| Sleep | baseline quiescent current between duty cycles |
| Wake | startup overhead before sensing or radio activity |
| Sample | on-sensor acquisition (e.g., motion, ADC, or other local sensing) |
| Listen | passive receiver or scanner active window (e.g., BLE scan burst) |
| Beacon | transmission event (e.g., BLE advertisement, packet, or uplink) |

A major use case for Biocosm is comparing policies under identical simulated worlds. For example, a fixed low-duty policy may conserve energy but miss short events, whereas an adaptive policy may increase capture after recent detections while undersampling isolated periods. Because ground truth is retained separately, Biocosm can quantify both efficiency gains and sampling bias.

### 4.6 Energy layer

The energy layer represents the current or charge cost associated with each state or action. Rather than assuming one hardware platform, Biocosm uses a configurable energy table. Users may supply theoretical estimates, datasheet-based values, bench-measured state currents, or empirically fit profiles.

Representative states follow the duty-cycle sequence shown in the policy layer of Figure 1:

This formulation supports hardware-agnostic comparison. A user can replace one energy profile with another without changing the biological world or analysis layer. The same simulated experiment can therefore compare not only policies, but also classes of instruments.

### 4.7 Analysis layer

The analysis layer evaluates observed logs relative to ground truth. Metrics may include event capture rate, false-positive rate, duration error, dyadic network error, energy per captured event, daily energy budget, estimated lifetime, and Pareto optimality across capture and energy.

A central output is the **capture–energy frontier** (Figure 4), where each candidate policy is represented by its energy cost and event capture performance. Policies on the frontier are those for which no alternative simultaneously improves capture and reduces energy. In multi-objective optimization terms, these are Pareto-optimal policies: no other policy can improve one objective without worsening at least one other (Fei Z et al. 2016; Mehta R 2020). This representation avoids treating any single policy as universally optimal and exposes trade-offs that researchers can choose from based on scientific goals, animal welfare constraints, battery capacity, and deployment duration.

**Figure 4:**
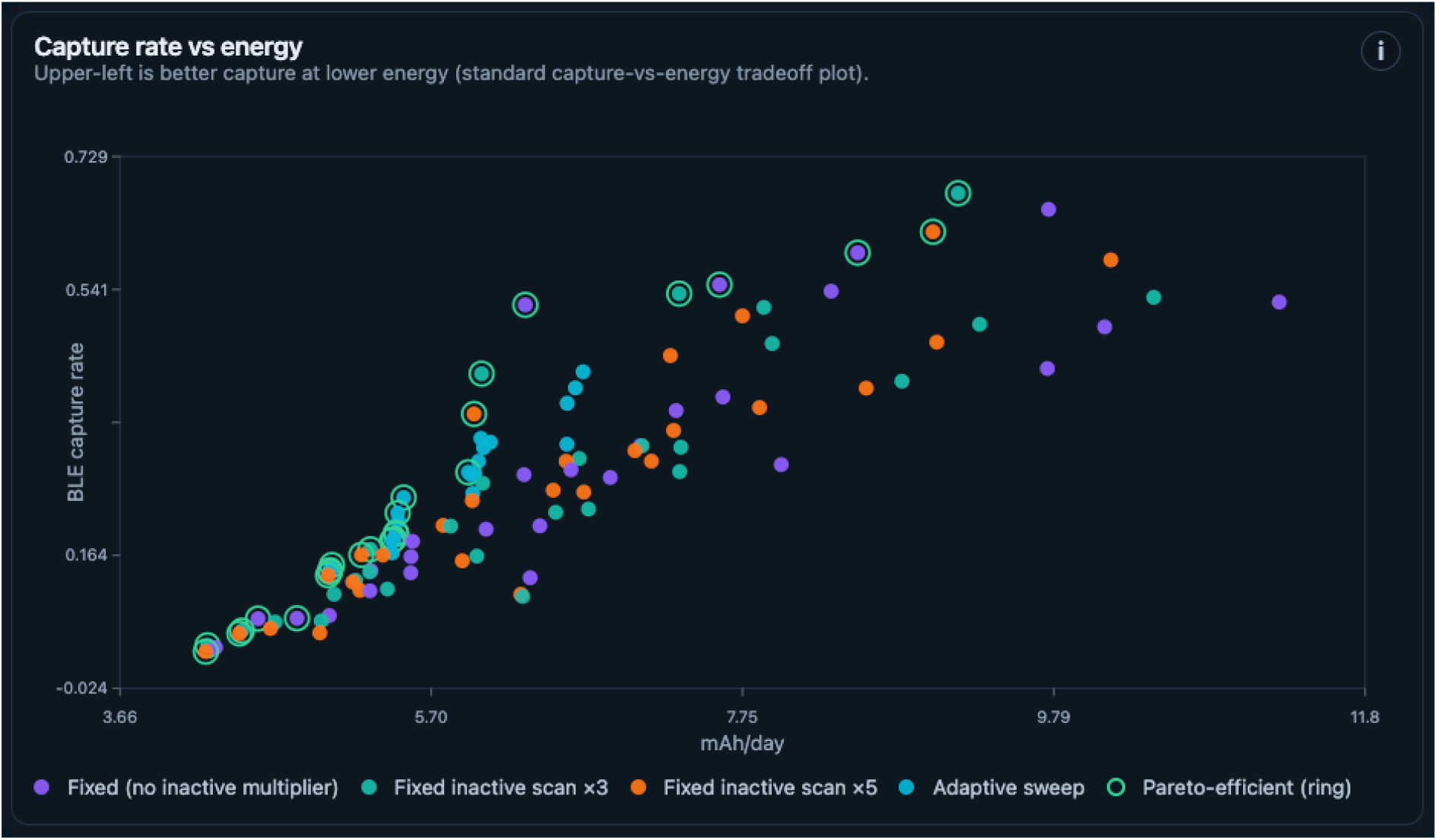
Capture rate versus daily energy for candidate sensing policies from a representative policy sweep (Pareto boundary; BLE reference implementation). Fixed schedules (with and without inactive-scan multipliers) and adaptive sweep policies are shown; green rings mark Pareto-efficient policies on the capture–energy frontier. Upper-left positions indicate higher capture at lower energy.

### 4.8 Software implementation

Biocosm is distributed as open-source, browser-based software. The live application is openly accessible at https://neurotech-hub.github.io/Biocosm/, with source code and build instructions at https://github.com/Neurotech-Hub/Biocosm. No server backend is required for core simulation, policy comparison, or export workflows; the application runs entirely in the client after loading from GitHub Pages or a local development build.

The implementation uses **React** and **TypeScript** for interface and core logic, **Vite** for bundling/development, an **HTML Canvas** renderer for enclosure visualization, and **Vitest** for tests. The simulation engine is organized into modular TypeScript components under `src/simulation/` that mirror the conceptual layers in Figure 1: ground-truth movement and circadian behavior (`world`), BLE dyad formation and RSSI-based detection (`radio`), deployable schedules including fixed-rate and motion/peer-adaptive policies (`policies`), epoch-level energy accounting (`energy`), and capture/efficiency metrics (`analysis`). Information flows in one direction (world → radio → policy → observed logs → analysis), and policies operate only on observable state; ground truth is retained for evaluation but is not exposed to the policy layer during simulation.

The web interface (Figure 2) groups controls for simulation timing, device/BLE parameters, and animal biology.

A *Build Simulation* action applies configuration changes before advancing the timeline. Epoch length, temporal resolution, communication schedules (BLE in the reference implementation), and the per-state energy table are all user-defined; visualizations and summary metrics are computed over the configured review window. Users can replay the entire run on the shared timeline—stepping through animal trajectories, true proximity events, scan/beacon activity, and logged captures—to confirm that the biological world and observation model produce plausible behavior before policy comparison or export.

In addition to single-run exploration, the application provides batch *Policy Sweep* grids over fixed and adaptive schedules and an in-app *Policy Optimizer* that fits surrogate response surfaces to sweep results and verifies candidate policies with full simulations. Workspace files support saving and reloading simulator configuration, view state, and sweep settings; outputs (species traits, event logs, and sweep summaries) can be exported as JSON or CSV for offline analysis.

Biocosm is not a packet-level Bluetooth protocol simulator. It abstracts firmware timing, scan/advertise scheduling, detection probability, and energy from user-specified duty cycles. The open-source application foregrounds a BLE proximity-logger workflow motivated by miniaturized wearables for free-living vertebrates (Gaidica and Dantzer 2022; Sanders et al. 2025), but burst durations, detection thresholds, and energy entries for sleep, wake, sample, listen, and beacon can be tuned to match datasheets, bench measurements, or alternate hardware. Example figures and sweeps in this manuscript use one documented reference configuration for reproducibility, not as prescribed settings for all users. Reproducible studies should report the application version, Git commit hash, random seed, species preset, energy model identifier, duty-cycle parameters, and sweep grid constants when publishing comparative results.

## 5 Example Analyses

The examples below are illustrative vignettes from documented reference runs (seed 42, *n* = 6 prairie voles, 24 h simulation, 30 × 20 m enclosure), not exhaustive benchmarks. They show the kinds of quantitative outputs Biocosm produces: per-animal time-series logs (JSON), and policy-sweep tables (summary and per-policy CSV). Together, these exports support replay, calibration checks, and offline comparison without rerunning the simulator.

### 5.1 Fixed versus adaptive sensing policies

A basic Biocosm experiment compares fixed and adaptive sensing policies in the same simulated world. The fixed policy observes at a constant interval throughout the deployment. The adaptive policy increases observation intensity after recent detections and returns to a lower duty cycle after a decay period.

In the reference balanced-adaptive run underlying Figure 3, the population spent a mean of 34% of epochs in motion-detected activity, logged 261 proximity detections over 24 h, and consumed 6.3 mAh per collar-day under the generic nRF52840 energy profile. Raw exports retain animal trajectories, true dyads, scan/beacon bursts, and cumulative energy minute-by-minute for inspection.

This example illustrates a central trade-off. Adaptive sensing may improve event capture per unit energy by concentrating effort during periods when additional events are more likely. However, this can create behavior-dependent sampling bias. Socially active periods may be oversampled, while isolated periods may be undersampled. Biocosm makes this bias measurable because the simulator retains both the latent event history and the observed logs.

### 5.2 Capture–energy frontier

A second analysis sweeps policy parameters to estimate a capture–energy frontier. Candidate policies vary observation interval, beacon interval, observation-window duration, adaptive gain, decay time, and energy-throttling thresholds. For each policy, Biocosm computes event-capture rate (BLE capture rate in the reference implementation) and daily energy cost under identical or replicated biological worlds.

Figure 4 summarizes one such sweep over 105 candidate policies in the same biological world.

Three illustrative takeaways: (i) the comparison baseline (balanced adaptive, 30 s scan / 10 s advertise) achieved 28% event capture at 6.3 mAh day^*−*1^; (ii) a competitive fixed schedule (10 s scan, 1 s window, 10 s advertise) reached 52% capture at similar energy (∼1.9× capture); and (iii) 27 policies lay on the Pareto frontier, spanning 4.2–11.3 mAh day^*−*1^ and 3–68% capture. High-duty fixed schedules can maximize capture (up to ∼68% in this grid) but at substantially higher energy.

The result is a design-space map rather than a single optimum: researchers can discard infeasible schedules (e.g., high capture above battery budget, or low energy with unacceptable miss rates) before deployment. Sweep exports include ranked summary statistics and per-policy raw metrics for reproducibility.

### 5.3 Observation bias in inferred social networks

A third analysis compares true and observed dyadic contact networks. The same ground-truth interaction matrix is passed through multiple sensing policies. Each policy produces an observed network, and Biocosm computes deviation from the true network.

This analysis is particularly important for social ethology. Two policies may have similar global event capture rates but different dyadic biases. This is consistent with animal-social sensing studies showing that measurement method and sampling design can change which encounters are detected, even when some aggregate network properties remain similar (Gelardi V et al. 2020; He P et al. 2023). For example, a policy that preferentially captures long contacts may preserve stable affiliations while missing brief exploratory interactions. A policy that adapts after detections may inflate repeated interactions among already-detected dyads. Biocosm allows these distortions to be quantified explicitly.

### 5.4 Hardware-agnostic energy profiles

A fourth analysis replaces the energy model while holding the biological world and policy logic constant. For example, one profile may represent an ultra-low-power beacon/listener, another a higher-sensitivity but higher-current sensor, and another a storage-heavy logger. This analysis demonstrates how Biocosm can compare classes of instrumentation without making the manuscript about a specific printed circuit board, microcontroller, or radio stack.

## 6 Validation and Calibration Strategy

Biocosm is intended to support multiple levels of empirical grounding. It does not require full hardware validation for theoretical exploration, but its predictive specificity increases as users add measured parameters.

This ladder clarifies the role of bench and field data. This staged calibration logic aligns with prior work calling for explicit device-performance assessment, validation in close-to-real conditions, and sampling-design analysis before strong biological inference is made (Boyland NK et al. 2013; He P et al. 2023; Ossi F 2016). Empirical current measurements are valuable, but they parameterize the energy model rather than redefine the paper as a device manuscript.

Similarly, field calibration improves the observation model without making the contribution a validation study of one collar architecture.

In practice, users can follow the ladder in Table 2: bench-parameterize energy and observation models, specify the biological world from species priors or pilot data, sweep policies in simulation to identify capture–energy frontier candidates, deploy only that subset, and compare field logs to held-out predictions. This sequencing narrows the design space before deployment and limits costly physical iteration while keeping calibration explicit.

**Table 2:** Calibration ladder for empirical grounding of Biocosm simulations.

| Level | Purpose and required data |
| --- | --- |
| Theoretical | Explore general trade-offs using plausible parameters and sensitivity ranges. |
| Bench-parameterized | Estimate energy and timing from bench measurements (state currents, transition costs, duty-cycle timing). |
| Field-calibrated | Match the observation model to deployment context using video, RFID, controlled-distance tests, synchronized loggers, or known encounters. |
| Held-out validation | Test predictive performance with separate calibration and validation datasets. |

## 7 Reproducibility and Implementation Considerations

A methods framework for ethological sensing should support reproducibility at the level of both biological scenarios and measurement policies. Biocosm therefore emphasizes seed-deterministic simulation, explicit parameter sets, exportable logs, and separable model layers (section 4.8). Reproducibility is especially important because policy comparisons are only meaningful if alternative policies are evaluated under equivalent biological conditions.

The framework should also preserve the distinction between simulation truth and deployable policy state. Firmware-like policies should operate only on information that would be available to a real device. Ground truth should be used only for evaluation. This constraint prevents optimistic policies that implicitly exploit impossible information.

Finally, Biocosm should make uncertainty visible. Observation models, energy models, and movement models should support sensitivity analysis. When parameters are uncertain, the appropriate output is not a single predicted lifetime or capture rate, but a range of plausible outcomes and a ranking of policies that is robust or fragile under parameter perturbation.

## 8 Discussion

Biocosm treats ethological instrumentation as an explicit measurement system. Data used for biological inference are produced jointly by animal behavior and instrument behavior. Animal proximity-logger validation studies show that recorded contacts can be shaped by distance, habitat, antenna orientation, body effects, device identity, and post-processing choices (Boyland NK et al. 2013; Drewe JA et al. 2012; Huels F 2025; Kirkpatrick et al. 2021; Levin II et al. 2015; Ossi F 2016; Prange S et al. 2006; Rutz C et al. 2015). Sampling-design studies in animal societies likewise show that sampling frequency, sampling duration, coverage, and measurement method can alter inferred interactions and network structure (Gelardi V et al. 2020; He P et al. 2023). A sensor that sleeps through a brief encounter does not merely save energy; it changes the dataset. An adaptive sensor that samples more aggressively after recent detections may improve capture, but it also changes the sampling design.

This policy-centered view connects ethological sensing to a broader engineering literature on duty-cycled wireless systems. BLE discovery models show that advertising interval, scan interval, and scan-window duration jointly determine discovery probability and latency (Cho K et al. 2015; Liu J, Chen C, and Ma Y 2012). Energy models of BLE further show that advertising, scanning, receiving, transmitting, sleeping, and state transitions contribute differently to total energy consumption (Kindt PH et al. 2020; Luo B and Liu J 2014; Siekkinen M et al. 2012). Asynchronous neighbor-discovery work generalizes this point beyond BLE by showing that low-power operation and active vigilance are fundamentally coupled through wake schedules and rendezvous latency (Dutta P and Culler D 2008; Sun W et al. 2014). Biocosm uses these ideas to treat observation policy as a scientific variable rather than a firmware detail.

The digital twin-inspired framing clarifies what Biocosm is, and what it is not. General digital-twin architectures describe virtual representations linked to physical systems through data flows, analytics, and application layers (Al-Ali AR et al. 2020; Khan LU et al. 2022; Minerva R, Lee GM, and Crespi N 2020). Medical and neurotechnology digital twins use related ideas to connect latent biological state, observation, prediction, and intervention (Corral-Acero J 2020; Popescu D 2024; Wang HE 2024). Biocosm adapts this architecture to ethological measurement as an executable measurement-process simulation: the latent biological world, observation process, sensing policy, energy model, and analysis layer are separable modules. This separation allows researchers to ask how alternative instruments would sample the same biological process.

The capture–energy frontier (Figure 4) is grounded in multi-objective optimization: Pareto-optimal solutions are those for which no objective can be improved without worsening at least one other (Fei Z et al. 2016; Mehta R 2020). For Biocosm, this means that a policy with higher event capture may not be preferable if it exceeds battery, mass, or deployment-duration constraints, while a low-energy policy may be unacceptable if it systematically distorts the behavior of interest.

The most important contribution of Biocosm is conceptual and methodological. Existing proximity-logging and biologging studies have shown the need for empirical calibration, context-specific validation, and caution when interpreting sensor-derived social data (Boyland NK et al. 2013; Drewe JA et al. 2012; Huels F 2025; Kirkpatrick et al. 2021; Levin II et al. 2015; Ossi F 2016; Rutz C et al. 2015). Biocosm does not replace those steps. Instead, it provides a reproducible environment for testing measurement assumptions before deployment and for interpreting how observation policies shape downstream inference.

The framework is intentionally hardware-agnostic. While wireless proximity sensing provides a concrete and scientifically important use case, the same structure applies to many energy-constrained modalities. Any device that must choose when to observe, transmit, compute, store, or sleep can be represented as a policy operating under an energy budget and an imperfect observation model.

## 9 Limitations

Biocosm does not eliminate empirical validation. Observation models are only as good as their parameters, and animal behavior models are simplifications. If the deployment environment differs substantially from the simulated world, predictions may be misleading. Bench-measured energy profiles may fail to extrapolate to extreme duty cycles, unusual temperatures, aging batteries, or unexpected firmware behavior. Wireless models that are useful for policy ranking may not capture packet-level channel dynamics, antenna detuning, enclosure effects, multipath, or collision behavior in dense deployments.

The framework also risks false precision if outputs are interpreted as exact predictions rather than model-dependent estimates. For this reason, Biocosm should be used for design-space exploration, sensitivity analysis, and policy comparison before deployment, not as a substitute for calibration or held-out validation.

A final limitation is that adaptive policies can create subtle sampling biases. Improving energy efficiency is not equivalent to improving scientific validity. Biocosm helps expose these biases, but it does not decide which bias is acceptable. That decision depends on the biological question.

## 10 Conclusion

As ethology, neuroscience, and bioengineering move toward long-duration, adaptive, and miniaturized sensing systems, the observation policy becomes part of the scientific method. Biocosm provides a measurement-process simulation environment for making that policy explicit, reproducible, and quantitatively comparable. By simulating the pathway from latent biological events to observed logs under energy constraints, Biocosm helps researchers evaluate sensing strategies before deployment and interpret the limitations of the data they collect.

